# How conspicuous are peacock eyespots and other colorful feathers in the eyes of mammalian predators?

**DOI:** 10.1101/514240

**Authors:** Suzanne Amador Kane, Yuchao Wang, Rui Fang, Yabin Lu, Roslyn Dakin

## Abstract

Feathers perceived by humans to be vividly colorful are often presumed to be equally conspicuous to other mammals, and thus to present an enhanced predation risk. However, many mammals that prey on adult birds have dichromatic visual systems with only two types of color-sensitive visual receptors (one sensitive to ultraviolet light), rather than the three characteristic of humans and four of most birds. Thus, understanding how these predators perceive color requires quantitative visual modeling. Here, we use a combination of reflectance spectroscopy, multispectral imaging, color vision modelling and visual texture analysis to compare the visual signals available to conspecifics and to mammalian predators for multicolored feathers from the Indian peacock (*Pavo cristatus*) as well as red and yellow parrot feathers; we also take into account the effects of distance-dependent blurring due to visual acuity. When viewed by tetrachromatic birds against a background of green vegetation, most of the feathers studied had color and brightness contrasts similar to values previously found for ripe fruit. By contrast, when viewed by dichromat mammalian predators, the color and brightness contrasts of these feathers were only weakly detectable and often did not reach detection thresholds for typical viewing distances. We furthermore show that the peacock’s erect train has undetectable color and brightness contrasts and visual textures when photographed against various foliage backgrounds. Given the similarity of photoreceptor sensitivities and feather reflectance properties across relevant species, these findings are consistent with many feathers of similar hue being inconspicuous, and in some cases potentially cryptic, in the eyes of their mammalian predators. These results suggest that many types of colorful feathers are likely to be cryptic to mammals while providing a communication channel perceptible to birds, while emphasizing the importance of understanding diverse sensory receivers in the evolution of animal coloration.

## Introduction

Ever since Darwin, colorful feathers such as the iridescent eyespots of the Indian peacock (*Pavo cristatus*) (Fig 1A) have been assumed to present salient visual signals readily detectable by their natural predators (Darwin, 1888; Ranjith and Jose, 2016; Ruxton et al., 2004). For this reason, these sexually-selected ornaments have been proposed to incur a cost due to increased predation. For example, as Zahavi stated in his paper introducing the handicap principle: “The more brilliant the plumes, the more conspicuous the male to predators” (Zahavi, 1975). Evidence for such countervailing selection pressures has been found in ornamented guppies preyed upon by fish (Endler, 1980) and in birds preyed upon by other birds (Møller and Nielsen, 2006). However, while this assumption is predicated on the predator being able to detect prey visual signals (Outomuro et al., 2017), no studies have tested whether this is true for the mammalian predators that prey on many birds. For example, the primary predators of adult peafowl are carnivorans (felids and canids, S1 Appendix), which all have dichromatic visual systems; i.e., they have only two types of cone visual receptors with distinct spectral sensitivities, not the four characteristic of most birds (Cronin et al., 2014) or the three found in most humans. More generally, felids (e.g., *Felis catus*) are a major threat to bird populations world-wide (Loss et al., 2015). Because dichromatic mammals lack red-green color discrimination, they are unlikely to detect many of the chromatic visual cues evident to birds and humans (Cronin et al., 2014; Miller and Murphy, 1995). Studies of visual ecology have considered how prey appear to various types of predators (birds, insects and fish) for many types of prey, including insects and birds (Håstad et al., 2005; Théry et al., 2005), fish (Endler, 1991), cuttlefish (Chiao et al., 2011), crustaceans (Nokelainen et al., 2017), primates (Sumner and Mollon, 2003) and lizards (Outomuro et al., 2017). Two previous studies also have studied the iridescence reflectance spectra of peacock eyespots and how they are perceived by peahens (females) (Dakin and Montgomerie, 2013; Loyau et al., 2007). As yet, no studies have compared how visual signals from peacocks and other avian prey appear in the vision of their mammalian predators.

**Fig 1.**
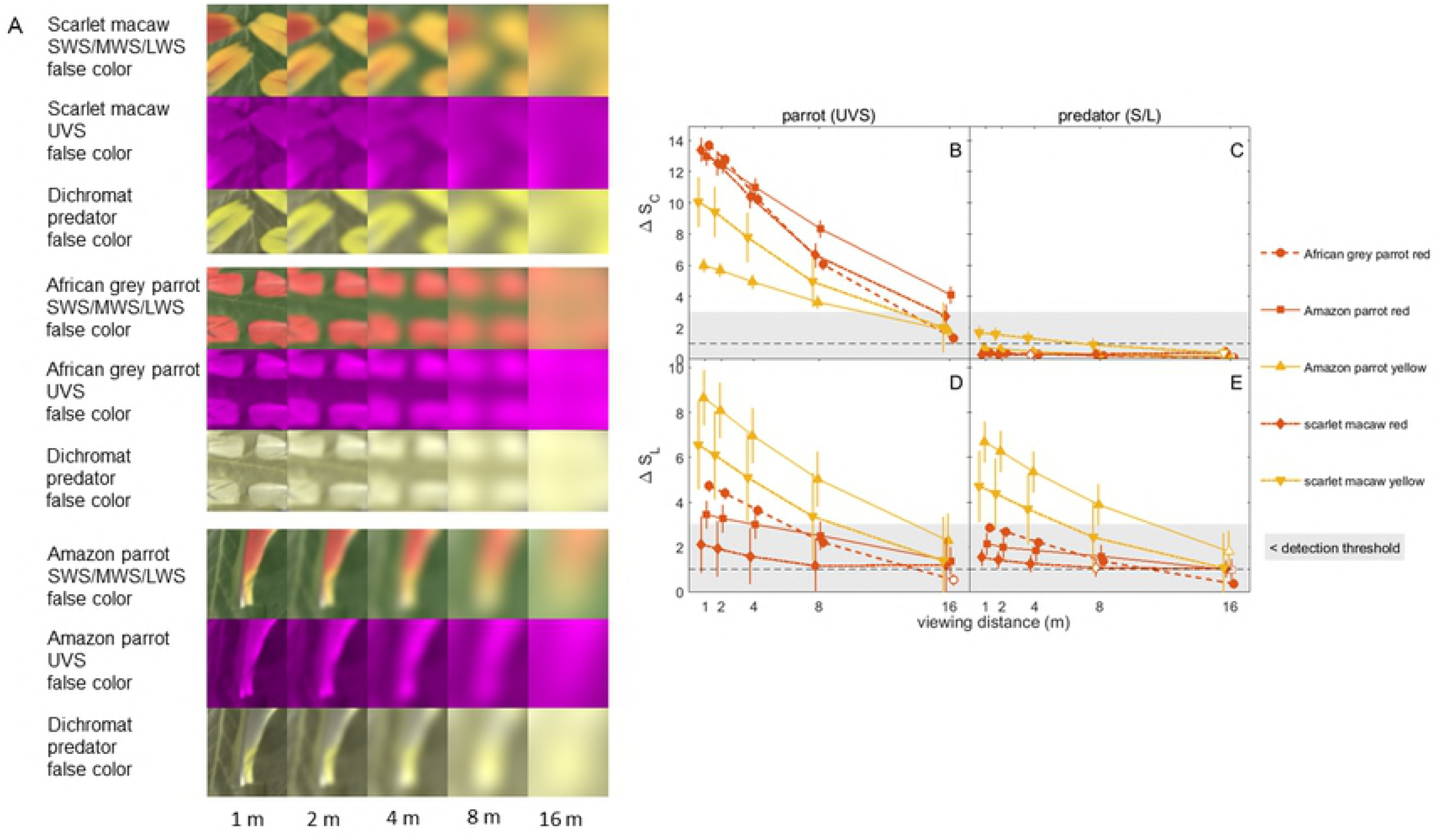
Peacocks and the model peacock train. A) An Indian peacock displaying his erect train to a peahen (female) in the foreground and B) another individual holding his train folded while walking. C) Model peacock train assembled from a collection of eyespot feathers used to evaluate the appearance of the train viewed against vegetation.

During courtship displays, male Indian peafowl (“peacocks”) attract mates by spreading, erecting and vibrating the fan-like train ornament (Fig. 1A), causing it to shimmer iridescently and emit mechanical sound (Dakin and Montgomerie, 2009; Dakin et al., 2016; Freeman and Hare, 2015). Several lines of evidence indicate that these feathers are assessed during mate choice: train-rattling performance by peacocks is obligatory for mating success (Dakin and Montgomerie, 2009), eye-tracking experiments have shown that train-rattling displays are effective at attracting and holding the peahen’s gaze (Yorzinski et al., 2013), and eyespot iridescence has been shown to account for approximately half of variation in male mating success (Dakin and Montgomerie, 2013; Loyau et al., 2007). Because peacocks spend the majority of their time in activities other than courtship displays even during the breeding season (Dakin and Montgomerie, 2009; Harikrishnan et al., 2010), any test of visual saliency must also consider the appearance of the folded train. Furthermore, because the peacock’s head, neck and breast are covered by iridescent blue contour feathers (Yoshioka and Kinoshita, 2002), the visual cues generated by this body plumage are also relevant for salience to potential mates and predators.

Here, we use multispectral imaging and reflectance spectroscopy to compare how detectable peacock feathers are to conspecifics and dichromatic mammalian predators (hereafter “dichromatic mammals”), as measured by color, brightness, and texture contrast relative to green background vegetation, following similar studies of prey that utilize camouflage against predators with a variety of visual systems (Stevens and Merilaita, 2011). Our goal was to test the assumption that colorful feathers that are highly conspicuous to conspecific birds are also readily detectable by these predators. To determine how generalizable our results were to other hues of colorful plumage, we also measured reflectance spectra and multispectral images of red and yellow parrot feathers. We then used psychophysical vision models to test whether conspecifics and dichromatic mammalian predators can readily detect the color and brightness contrasts between feathers and green vegetation. Our analysis modeled the appearance of feathers at various distances to determine when each observing species could distinguish color patches relative to the surrounding environment.

In addition to color cues, visual salience depends on the presence of pattern features that are perceptually discriminable from the background. To determine whether predators might detect the peacock’s train using such visual texture cues, we analyzed images of the model train relative to that of background vegetation using two pattern analysis methods motivated by visual processing in vertebrates (Stoddard and Stevens, 2010). Granularity analysis is a spatial filtering method that determines the contributions to image contrast of features with different sizes; this image processing technique has been used to compare pattern textures in studies of cephalopod, avian egg, fish and shore crab camouflage, as well as humans searching for objects against various backgrounds (Akkaynak et al., 2017; Barbosa et al., 2008; Nokelainen et al., 2017; Stoddard and Stevens, 2010; Troscianko et al., 2017). A second method, edge detection, provides a complementary measure of texture complexity by using image processing to detect sharp gradients in intensity (Stoddard et al., 2016).

## Materials and methods

### Feather samples

Five Indian peafowl eyespot (Fig 2A), three blue peacock contour breast feathers (Fig 2B), four scarlet macaw (*Ara macao*) wing feathers (two red and six yellow patches total) (Fig 3A), two Amazon parrot wing feathers (two red and two yellow patches total) (Fig 3B), and four red African grey parrot (*Psittacus erithacus*) tail feathers (Fig 3C) were obtained from Moonlight Feather (Ventura, CA USA) and Siskiyou Aviary (Ashland, OR USA). Because the psittacofulvin pigments in parrot feathers have reflectance spectra with similar spectral features to red and yellow carotenoid pigments (Shawkey and Hill, 2005; Toral et al., 2008), our results should be representative of red and yellow feathers in general. Our number of replicates for the peacock eyespots agree well with recommendations from a study of intraspecies variations in feather color measures (Dalrymple et al., 2015); however, we were limited by availability to fewer replicates for the parrot feathers. For mounting, eyespot feathers were cut off below the outermost colored ring at the proximal end. All feather types were mounted on black matte art quality paper with a magnetic backing that adhered to the tilt stages used for spectroscopy and multispectral imaging. Feather samples were stored without compression in sealed boxes in acid-free envelopes at 75% relative humidity and ambient temperature (22 ± 2 deg C). The different peacock eyespot color patches (colored rings and central disk) are referred to using the names and two letter abbreviations indicated in Fig 2A.

**Fig 2.**
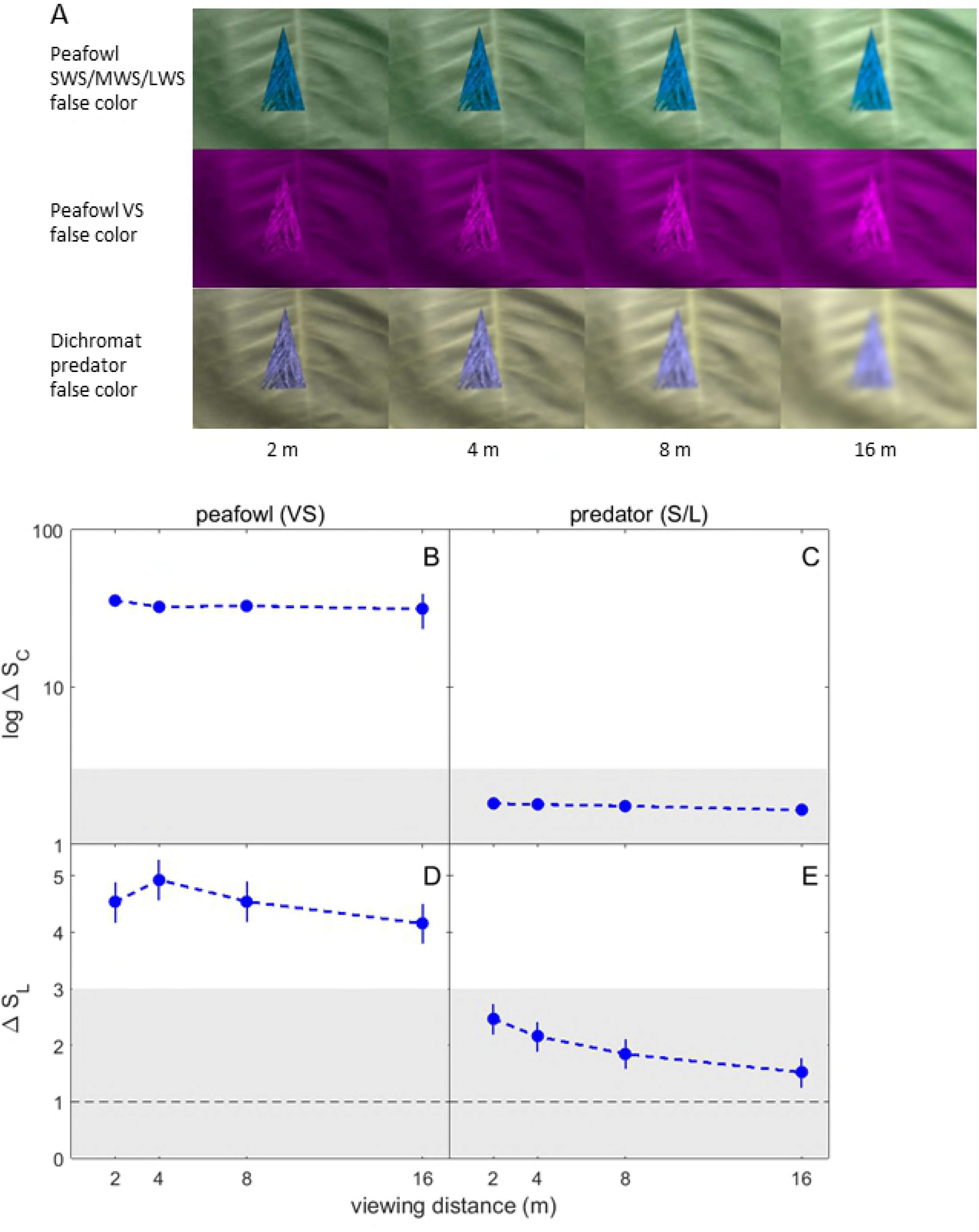
Peafowl cone sensitivity spectra and peacock feather vs green vegetation reflectance spectra. (A) An Indian peacock eyespot feather showing the color patch names used in the analysis. (B) Peacock blue breast plumage. (C) Comparison of the cone photoreceptor spectral sensitivities for the Indian peafowl and ferret, which has dichromatic color vision very similar to that of cats and dogs. All spectra are multiplied by the D65 illuminance spectrum used to model sunlight and normalized to unit area. Reflectance spectra of (D) peacock feather eyespots and (E) peacock iridescent blue body plumage and the green saucer magnolia (*Magnolia x soulangeana*) leaf used as a background for the feather sample images.

**Fig 3.**
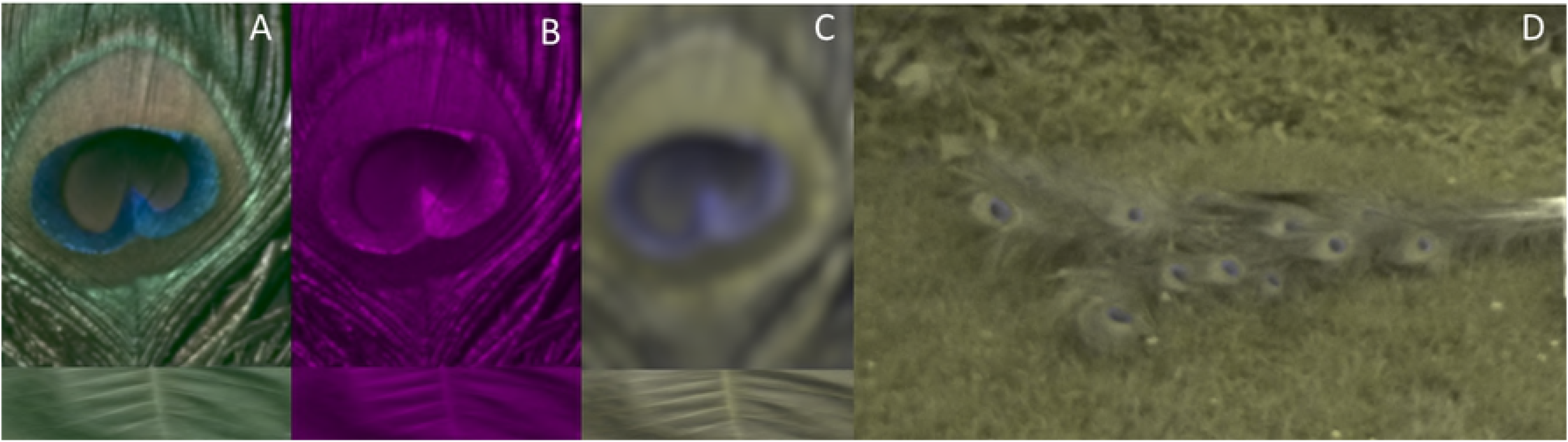
Parrot feather images, cone sensitivity spectra and feather reflectance spectra. (A) Scarlet macaw, (B) Amazonian parrot and (C) African grey parrot feather samples. (D) Comparison of the cone photoreceptor spectral sensitivities for the blue tit, which has tetrachromatic ultraviolet sensitive (UVS) color vision similar to that of parrots, and the ferret, which has dichromatic color vision similar to that of cats and dogs. All spectra are multiplied by the D65 illuminance spectrum used to model sunlight and normalized to unit area. (E) Reflectance spectra of the parrot feather red and yellow patches studied here.

We also assembled an array of 28 peacock feathers (Fig 1C) to create a model train arranged to match the geometry of eyespots in actual peacock trains ((Dakin and Montgomerie, 2013)); this was used to simulate the appearance of the train during display (when the train is erect) or during walking, perching or standing (when the train is held horizontally; see Fig 1B). In their native range in India and Pakistan, peafowl are reported to live in a variety of habitats, including open moist and dry-deciduous forest, scrub jungle, and adjacent grasslands, and their breeding season is reported to coincide with the start of the rainy season (Gokula, 2015), after which eyespot feathers are shed by molting (Beebe, 1918; Sharma, 1974). We used as background foliage for feather and model train images various plants (grass, brush and trees) native to the northeast USA (S2 Appendix). Because green flora have generic reflection spectra due to chlorophyll absorption (Jensen, 2009), the plants used in this study should be representative of the color and luminance of those found in the native environments of peafowl and many other bird species.

### Vision models

The Indian peafowl’s visual system has four classes of color-sensitive (chromatic) single cone cells: violet (VS), short (SWS), medium (MWS) and long (LWS) wavelength-sensitive cones, and one type of double cone that is sensitive to brightness (luminance) (Hart, 2002). In order to illustrate their spectral responses under natural illumination, Fig 2C shows the peafowl cone’s spectral sensitivities *S_r_(λ)* for the r^th^ photoreceptor type (including ocular media and oil droplet transmission) multiplied by the CIE D65 irradiance spectrum, *I(λ)*, and normalized to unit area; we used this standard illuminant because of its close match the solar irradiance spectrum for the elevation angles found for actual peacock displays (Dakin and Montgomerie, 2009; Spitschan et al., 2016). To model the tetrachromatic UVS (ultraviolet-sensitive) vision of parrots we used blue tit (*Cyanistes caeruleus*) cone spectral sensitivities (Hart et al., 2000; Troscianko and Stevens, 2015). (Fig 3D).

The visual systems of dichromatic mammalian predators have been studied for a variety of genera, and found to include S (blue-to near-UV-sensitive) (Douglas and Jeffery, 2014) and L (green-sensitive) cone populations in all carnivorans studied to date, including felids (Guenther and Zrenner, 1993) and canids (Jacobs et al., 1993). Behavioral studies have confirmed that domestic cats (Clark and Clark, 2016) and dogs (Kasparson et al., 2013; Neitz et al., 1989) have dichromatic color vision. Brightness signals in dichromatic mammals are assumed to be due to only the L cones (Osorio and Vorobyev, 2005). We used ferret (*Mustela putorius*) cone spectra (Troscianko and Stevens, 2015) to model dichromat vision because ferret spectral peaks agree closely with those of cats and dogs (i.e., ≤ 4.4% for S and ≤ 1.4% for L cones) (Calderone and Jacobs, 2003; Guenther and Zrenner, 1993; Jacobs et al., 1993) (Fig 2C). While many carnivorans are primarily nocturnal or crepuscular, at low light levels, photopic chromatic signals will be weak and visual signals will be dominated by luminance contrast via rods or double cones. Thus, we consider high luminance photopic conditions as the best case scenario for visual detection by these predators.

### Multispectral imaging

Multispectral images were recorded using a GoPro Hero 4 Silver Edition camcorder (GoPro Inc, San Mateo, CA USA) modified for full spectral imaging by replacing its original lens and infrared (IR) filter with a quartz lens transparent to < 300 nm (Igoe et al., 2013; Prutchi, 2016). Because the spectral response of this camera’s IMX117 Exmor-R CMOS sensor (Sony Corp., Tokyo, Japan) is sensitive throughout the visible and near-UV, these cameras have been used in multispectral imaging (Vogt and Vogt, 2016; Yun et al., 2016) (S1 Fig). Multispectral photographs were recorded at 3000 × 2250 pixel resolution and the GoPro settings medium field of view, Protune CAM-RAW mode (for no white balance compensation), flat color, low sharpness, ISO 400, exposure -2, night mode (to enable shutter speed control), auto shutter and spot meter on. Each sample was photographed for each geometry and illumination condition to give two multispectral images: 1) a UV image using an Andrea-UV filter (< 1% transmission for > 400 nm; UVIROptics, Eugene, Oregon USA; 2) a visible RGB (red, green, blue) image using two UV-IR cut filters to pass 400-700 nm light (Hoya Corp., Tokyo Japan). Filter transmission spectra were measured using the methods described below in “Reflectance Spectroscopy” (S1 Fig). The camera’s large depth-of-field eliminated the need for refocusing between visible and UV images. To maintain constant camera alignment between photographs, we mounted the camera rigidly using optical mounts (Thorlabs, Newton NJ, USA) and attached filters using quick-release Xume magnetic adapters (Panalpina Inc., Port Reading, NJ, USA); all images were taken using a remote trigger. Each feather image included a model Micro FSS08 8-step grayscale diffuse reflectance standard (Avian Technologies, New London, NH USA) mounted level with the sample plane for calibrating absolute reflectance (Troscianko and Stevens, 2015). Images of the model train included a larger 6-step grayscale and color checker chart (DGK Color Tools WDKK Waterproof, Digital Image Flow, Boston MA USA). Reflectance spectra for each grayscale in each filter and camera color channel combination were measured using the methods described below in “Reflectance Spectroscopy”. Each image also included an object of known size for spatial calibration.

All samples were mounted on a tripod for imaging (S2 Fig). Three sets of multispectral images each were obtained with the model train held erect and held horizontal viewed from the side. Peacock eyespots were oriented with their rachis vertical to simulate their average orientation in the erect train during courtship displays and the model train was oriented in a variety of directions to simulate the variation in appearance of the iridescent train eyespot feathers during courtship display, standing and walking. The camera was mounted on a second tripod a distance 20.0 ± 1.0 cm from feather samples and 1.70 to 2.00 ± 0.05 m from the model train. For feather samples, the camera was oriented to record images at normal observation angle (θ = 0 ± 2 deg) with respect to the feather sample plane (Fig 4). The size of feather sample images was 55 mm x 67 mm, corresponding to 7.3 pixel/mm. Images were captured during June-July 2018 in the Haverford College Arboretum (latitude, longitude: 40.0093° N, 75.3057° W) for 24.2 ± 0.2 deg C and 55.5 ± 1.5 % relative humidity. All feather samples were illuminated by direct sunlight with an azimuthal angle Ψ = 45 ± 3 deg clockwise from the camera’s optical axis and at solar elevation angles Φ = 30 ± 3 deg, corresponding to an angle α = 52 ± 3 deg between the observation and illumination directions (Fig 4). These illumination and observation angles agree with those measured for female peafowl observing courtship displays (Dakin and Montgomerie, 2009). In general, these solar angles hold for the early morning times when most birds are most active (Robbins, 1981). Optimal color contrasts for non-iridescent feathers have been found to correspond to the range of observation-illumination angles α used in this study (Barreira et al., 2016); this is relevant because pigment-based colors can appear in combination with structural coloration (Shawkey and Hill, 2005). Furthermore, for this observation geometry, the bird’s body subtends the greatest visual angle. The peacock eyespot feather samples were surrounded by additional loose green barbs to simulate their setting in the actual train, while the parrot feather samples were surrounded by a saucer magnolia leaves picked ≤ 1 hour before image capture. We also imaged a variety of green leaves for comparison (S3 Fig). Black velvet fabric was mounted behind the feather samples to limit backscattered light and a lens hood was used to reduce lens flare. The model peacock train was photographed against a variety of foliage backgrounds for solar elevation angle between 37 to 55 deg.

**Fig 4.**
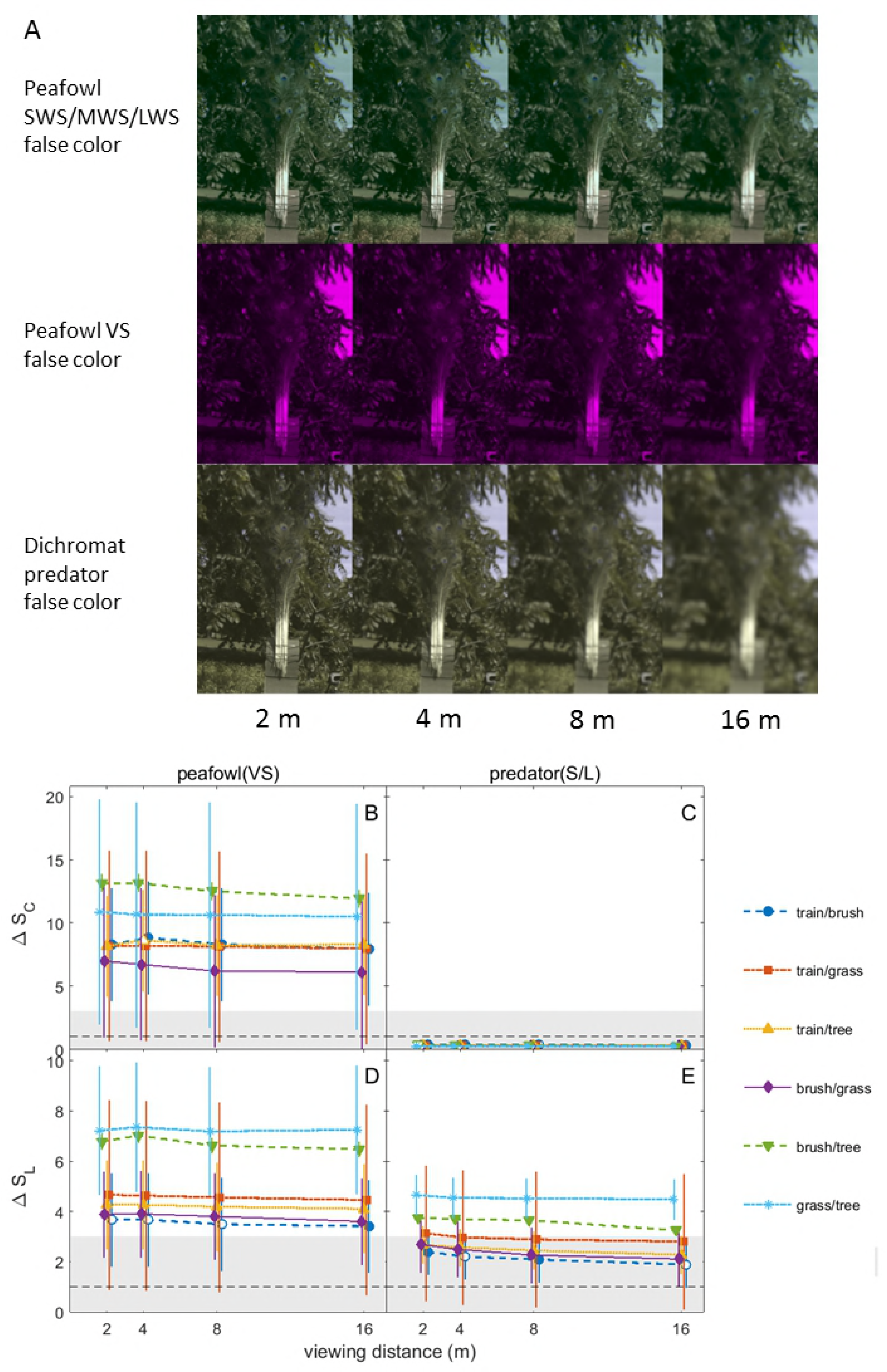
Multispectral imaging geometry showing the angles of observation and illumination. Multispectral images were first processed using custom scripts written in MATLAB v15a with the Machine Vision, Signal Processing and Fitting toolboxes (MathWorks, Natick MA USA); all code is available on figshare at https://figshare.com/s/f8add694af9c79de7f76. Images stored as jpeg files were calibrated and corrected for lens distortions using the MATLAB Camera Calibration application, and then corrected for perspective distortions using MATLAB’s *fitgeotrans* and *imwarp* commands. Images captured using the UV and visible filters were checked for alignment by hand and then converted into linearized and normalized measures of reflectance, as explained under “Quantitative visual signal analysis” below.

To account for distance-dependent blurring due to each viewing species’ visual acuity (Barnett et al., 2018; Caves and Johnsen, 2018), multispectral images with linearized intensities were spatially filtered before analysis to model the effect of viewing distance on contrasts between feathers and background foliage, and its effect on contrasts within the patterned eyespot feathers (See details in S3 Appendix). While peahens view peacock courtship displays at nearby distances ≥ 1 to 2 m (Dakin and Montgomerie, 2009), we also modeled a variety of greater viewing distances (2, 4, 8 and 16 m). Color patches were defined by hand in the original images and used for each modeled distance for uniformity. After spatial filtering and before color and brightness analysis, we sampled intensity values in the multispectral images on a square grid with spacing equal to a visual acuity disk, following (Endler, 2012). To model the effect of spatial filtering on the peacock’s blue head, neck and breast plumage, we used an image with green foliage background with an approximately peacock-shaped cutout of the blue plumage superimposed; spatial filtering was performed using peacock body dimensions (Talha et al., 2018) to define the composite image’s effective spatial scale.

To approximate how the feathers appear in each viewer’s visual system, three type of false colo images (ultraviolet, human visible RGB and viewer false color) were created from the multispectral images in MATLAB using square-root transformed cone quantum catches, *Q*_*pr*_ normalized to the maximum value of the brightest cone quantum catch on each image. To represent the tetrachromatic vision of peafowl, an RGB image was created from the computed LWS, MWS and SWS data, respectively and a magenta image was created using the VS cone data. To model dichromatic mammalian predator vision, we made up a single false color imag using blue to represent the S cone and yellow to represent the L cone quantum catch values.

### Reflectance spectroscopy

We measured reflectance spectra using a model USB2000+ spectrometer and OceanView software (Ocean Optics, Largo FL, USA) over the wavelength range 300-850 nm, using 100 m integration time, 3 pixel boxcar averaging (corresponding to the optical resolution of 6.5 pixel = 2.06 nm FWHM), and averaging over 5 samples. All spectra were recorded in a dark room. Samples were illuminated by an Ocean Optics PX-2 Pulsed Xenon Light source triggered at 20 Hz using square wave pulses from a model 330120A function generator (Agilent Technologies Wilmington, DE, USA); the source was turned on and allowed to warm up and stabilize for 15 minutes before data collection. Light for illumination and detection was carried in P400-1-UV· VIS optical fibers transparent to 200 nm (Ocean Optics). We used two PTFE white standards with flat 99.0% reflectance over 300-700 nm: a Spectralon USRS-99-010-EPV (Labsphere, North Sutton, NH USA) and a model SM05CP2C (Thorlabs). White standard and dark current were measured every fifteen minutes. For each feather and each measurement geometry, raw reflectance spectral data were recorded for each feather color patch sample radiance, AR, white standard radiance, AR_r_ and dark current, D. The reflectance spectrum, 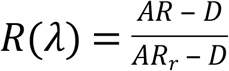 was smoothed over a wavelength interval of 20 nm using Savitzky-Golay smoothing in Origin; this reduced high frequency noise but did not change reproducible features of the spectra peak shapes.

Transmission spectra for the filters used in multispectral imaging were measured by recording the spectrum of light reflected from the white standard with and without the filter inserted into the light path with its face at normal incidence to the incident light. Reflectance values for color and gray standards were measured using a RPH-SMA reflectance probe stand (Thorlabs, Newton NJ USA) with the illuminating light at 45 deg to normal incidence and detected at normal incidence. The reflectance goniometer for feather measurements used (S2 Fig) was adapted from previously published designs (Van Wijk et al., 2016) (S2 Fig) but with an additional angular degree of freedom to allow measurement of the bidirectional reflectance distribution function, in which the angle of observation and illumination are not confined to the specular reflection geometry (Vukusic and Stavenga, 2009). Both the illumination and detection optical pathways were focused using a 74-UV lens (Ocean Optics) to a 2 mm diameter spot at about 5 cm from the output surface of the lens. The feather samples were realigned every time the angle of illumination and/or detection was adjusted to ensure both beams focused on the same region of the feather. To assess reproducibility of spectra for the same color patch on each feather, we measured each set of spectra three times for each sample after dismounting and remounting each sample.

### Quantitative visual signal analysis

We computed the color contrast, *Δ*S_c_, between color patches in the feathers and background vegetation in our multispectral images using the receptor noise limited color opponent model, which has been shown to predict behavioral thresholds for visual signals in birds, humans and insects (Vorobyev and Osorio, 1998). All calculations were performed using a custom MATLAB script, which was tested by verifying that it computed the same values as the multispectral analysis software package MICA version 1.22 (Troscianko and Stevens, 2015). First, intensity values, *V*, from each multispectral image were corresponded to the actual reflected irradiance, R, for this camera by an S-log transformation:

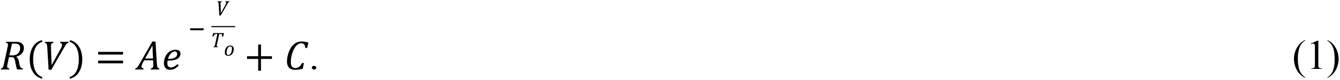

The parameters *A, T_o_* and *C* were obtained from nonlinear least squares fits in MATLAB (adjusted-R^2^ ≥ 0.997) of the measured *V* and *R* values for each pixel in each RGB channel of the image. The resulting fits then were used to convert measured intensity values for each p^th^ color patch into linearized and normalized reflected intensities (range [0,1]) for each combination of filter and RGB image channel. To compute the color and brightness contrasts, these intensities were converted into the cone quantum catch values, *Q_pr_*, for each of the viewer’s r^th^ cone photoreceptors:

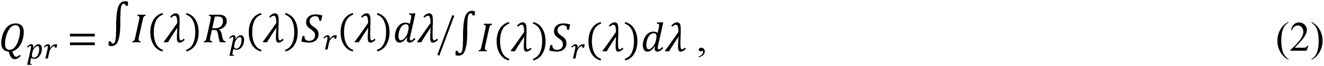

where *I(λ)* is the illumination spectrum, *S_r_(λ)* is the r^th^ cone receptor’s normalized spectral sensitivity and *R_p_(λ)* is the pth patch’s reflectance spectrum. Because birds and mammals are known to achieve color constancy under a wide variety of illumination conditions (Kelber and Osorio, 2010; Olsson et al., 2016), this equation also incorporates the von Kries transformation, a mechanism for maintaining color constancy (Stoddard and Prum, 2008). To accomplish this conversion, we used MICA to compute the parameters of a polynomial cone mapping between the UV blue channel and the visible RGB channels of the multispectral images recorded by our filter-camera system and the corresponding cone quantum catches, *Q_pr_* (Stevens et al., 2007).

This software finds the optimal mapping using our measured filter transmission and camera RGB spectral response curves with either the dichromatic ferret or tetrachromatic peafowl cone spectral sensitivities, the CIE D65 illumination spectrum and a large database of natural spectra. The net effect is to combine all measured values of linearized and normalized reflectance to compute the quantum catch, *Q_pr_*, of each r^th^ cone (r = S or L for dichromats and r = VS, SWS, MWS, or LWS for tetrachromats) for the p^th^ sample color patch. Using a linear 2-way interaction cone mapping model, we obtained a near perfect fit for each visual system: ferret (R^2^ ≥ 0.999), peafowl (R^2^ ≥ 0.996) and blue tit (≥ 0.990 UVS cone, ≥ 0.998 all other cones).

The resulting cone quantum catch values, *Q_pr_*, can be used to compute normalized color space coordinates, for the pt^h^ color patch: 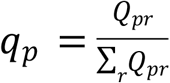 For tetrachromats, the receptor index *r* = VS or UVS, SWS, MWS, LWS and q_p_ = (v,s,m,l), while for dichromats r = S, L and q_p_ = (sw,lw). After normalization, this corresponds to a three-dimensional tetrachromat color space for birds and a one-dimensional colorspace for dichromats, here chosen to rely on sw. To validate the results of our multispectral imaging code, we compared dichromat color space sw coordinates computed by both MICA and our MATLAB code (sw_M_) from our multispectral images with those computed directly from reflectance spectra (sw_R_) for six color chart squares. Use of the camera and UV/visible filter cone mapping model was validated for multispectral image analysis by the goodness of the linear fit, zero intercept and unit slope, between the two sets of color space measures gave sw_M_ = (0.018 ± 0.031) + (1.00 ± 0.07) × sw_R_ + (adjusted-R^2^ = 0.993) (S1 Fig).

To compute color contrasts, *Δ*S_c_, we first computed the r^th^ cone’s log-linear quantum catch (Weber-Fechner), log *Q_rp_*, for each p^th^ patch. This was used to compute the difference in r^th^ cone response for the pq^th^ patch pair, Δ*_rpq_ =* log*Q_rp_* - log*Q_rq_.* The color contrast then is computed from differences between opponent cone pairs weighted by receptor noise. Dichromats have only S/L receptor opponency, so for them, 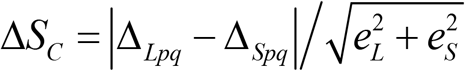 (Vorobyev and Osorio, 1998). The corresponding equation for color contrast in tetrachromats is more complicated because all six possible combinations of the four single cones pairs should be considered (Kelber, 2016):

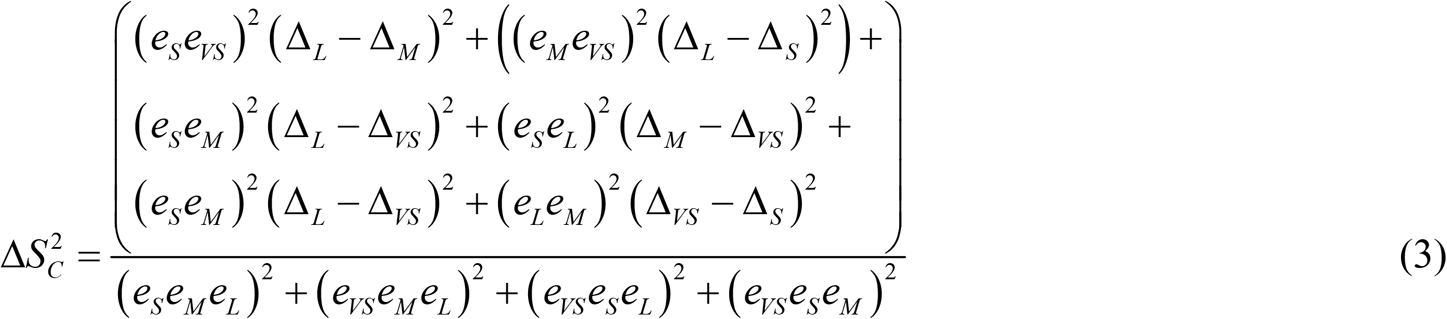

For bright illumination levels, receptor noise is assumed to be a constant determined only by the Weber fraction, *w_f_* and the relative population density, *g_r_*, for each r^th^ cone class (Renoult et al., 2015): 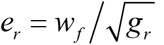. For peafowl, we used the value for chromatic Weber fractions of *w_f_*= 0.06 for L cones for domestic chickens based on color discrimination (Olsson et al., 2015). Receptor noise values for the other single cone classes were estimated using mean peafowl relative population densities *g_r_* = (0.477,0.892, 1.047, 1) for (VS, SWS, MWS, LWS) (Hart, 2001), yielding *e_r_* = (0.087, 0.064, 0.06, 0.06). For parrots, we used g_r_ = 0.25:0.33:1.05:1 and *W_f_*= 0.105 found for spectral sensitivity in *Melopsittacus undulatus* (Lind et al., 2014), corresponding to *e_r_* = (0.210, 0.182, 0.102, 0.105). Because color discrimination has not been measured for other mammals (Olsson et al., 2017), following (Stoddard et al., 2019) we used *W_f_*= 0.22 found for brightness discrimination in domestic dogs (range 0.22-0.27) (Pretterer et al., 2004). The relative cone population fractional densities measured for domestic cats (Linberg et al., 2001) give a mean g_r_(S,L) = (0.12,1); similar ratios have been reported for various wild felids (Ahnelt et al., 2006) and domestic dogs (Mowat et al., 2008). This gives the estimated predator receptor noise for color discrimination as (*e_S_, e_L_*) = (0.64, 0.22).

The brightness contrast, *Δ*S_L_, between each pq^th^ pair of color patches was computed from the quantum catches, *Q_Lp_* for the p^th^ color patch for the spectral response for the luminance channel (double cones for birds and L cones for dichromat predators) using *Δ*S_L_ = (log Q_Lp_ - log Q_Lq_)/w_f_, where *W_f_* is the Weber fraction for brightness discrimination. For birds, we used *w_f_* = 0.18 measured for double cones in budgerigars (*Melopsittacus undulatus*) (Lind et al., 2013); for comparison, lower values 0.10 have been found for pigeons (Hodos et al., 1985) and higher values ≥ 0.24 for chicks of the domestic chicken (Jones and Osorio, 2004). For predators, we used *w_f_* = 0.22 for brightness discrimination in domestic dogs as explained above; for comparison, *w_f_* = 0.10 in humans, and *w_f_* ≤ 0.45 in other mammals (Maertens and Wichmann, 2013; Olsson et al., 2017).

Color and brightness contrasts are interpreted in units of just noticeable distances (JND), with JND = 1 corresponding to the threshold for two patches to be discriminable under ideal illumination and viewing conditions when suitable data exist for the visual system being modeled (Olsson et al., 2015; Vorobyev and Osorio, 1998). Behavioral studies have shown that birds detect colorful fruit at a rate that correlates with increasing color (but not brightness) contrast for values >> 1 JND (Cazetta et al., 2009), while in lizards, the probability of discriminating a color from its background was found to be < 20% at 1 JND and to scale approximately linearly over the range 1 ≤ JND ≤ 12 (Fleishman et al., 2016). Behavioral tests in zebra finches have found that color contrast detection thresholds range from JND = 1 to 2.5 to 3.2, depending on background color (Lind, 2016). Following (Siddiqi et al., 2004), we therefore assume that the contrast detection threshold is approximately JND = 1 and we define contrasts in the range 1 < JND ≤ 3 as weakly detectable.

### Pattern Analysis

To model the perception of visual texture of the peacock’s train viewed against foliage, we also performed granularity pattern analysis on the model peacock train photographs, using MATLAB code adapted from (Barbosa et al., 2008) and MICA’s granularity texture analysis package. In granularity analysis, an image based on the luminance channel is filtered using an FFT bandpass filter centered at a series of spatial frequencies (granularity bands). For each bandpass-filtered image, the “pattern energy” (a measure of information at each spatial scale) is computed as the standard deviation of its pixel intensity values. The “granularity spectrum” then is defined as pattern energy vs granularity band. Granularity analysis was performed on the model peacock train images processed for the dichromatic predator luminance channel as explained above. Granularity spectra were computed for polygonal regions of interest (ROI) encompassing the entire model train and each type of surrounding vegetation (i.e., tall grass, brush or trees). To compensate for the effect of ROI shape and background, we used the following method adapted from MICA. For each ROI, we first computed a masked image in which all regions outside the

ROI were replaced by a black background. Next, we created a mean masked image in which the region inside the ROI in the masked image was replaced by the mean intensity within the ROI. Identical granularity calculations were performed on both images and their difference was used to create a shape-independent granularity spectrum.

We also computed summary statistics for comparing textures of the model train and its background: total energy (the energy summed across all filter bands, which increases as pattern contrast increases), peak filter size (the granularity band at peak energy; larger peak filter size corresponds to smaller most prevalent feature size), and proportion energy (the maximum energy divided by the total energy, a measure of how much of the spectral energy lies at the most prevalent feature size; this decreases with increasing pattern scale diversity). Granularity spectrum were plotted as “normalized energy” (pattern energy divided by total energy) vs granularity band to give a measure of how pattern information is distributed across spatial scales. Images with a uniform distribution of pattern scales have correspondingly uniform granularity spectra, while images dominated by a single feature scale should have strongly peaked spectra.

Edge detection of the luminance channel image provides an alternative measure of pattern complexity. We used the Canny edge filter in MATLAB to find edges using sigma = 3 and threshold = 0.15 to 0.20 (relative to maximum luminance image intensity set to 1) (Lovell et al., 2013; Melin et al., 2016). Model train images were first log transformed and then processed using contrast-limited adaptive histogram equalization (Zuiderveld, 1994) (*adapthisteq* in MATLAB) to detect texture edges in regions of widely differing illumination. The edge fraction (percentage of edge pixels in each ROI) was then used to compare the model train with various types of vegetation in the background (Stoddard et al., 2016); higher edge fractions indicate a more complex pattern with more spatial features.

### Statistics

Our analysis of color and brightness contrasts followed the two-step process recommended in (Maia and White, 2018). First, to determine whether the mean color and brightness contrasts between each patch pair had a statistically significant difference given their variances, we used PERMANOVA modified for non-normal, heterogeneous data (Anderson et al., 2017) implemented in the software package FATHOM (Jones, 2017) using 1000 bootstrap samples. Note that is it possible for this difference to be statistically significant, but to have value too small for it to be perceptually distinguishable. We therefore determined the effect size (how perceptually distinct each color patch pair) as follows. We first drew with replacement 1000 bootstrapped sample pairs using the MATLAB command *datasample*, and computed the mean *Δ*S_C_ and *Δ*S_L_ for this bootstrap resample. These mean contrasts were averaged over all images to get the mean and s.e.m. for each color patch pair for each sample; the grand mean and s.e.m. then was computed by averaging over all replicates for each feather type. Grand means and s.e.m. for the texture summary statistics were calculated from the mean of each statistic taken over all model train data for the train, grass, brush, and tree foliage. All results are reported as grand mean [95% CI = 2 s.e.m].

### Data Accessibility

All data and software required in order to replicate all of our results are archived either in the supplemental materials or at https://figshare.com/s/f8add694af9c79de7f76.

## Results

### Reflectance spectroscopy

Reflectance spectra for peacock eyespot feather color patches (Fig 2D) and peacock iridescent blue plumage (Fig. 2E) had spectral peaks consistent across repeated measured to 4 to 16 nm (95% CI) and exhibited similar spectral peaks and overall shape to those measured for zero elevation angle (Dakin and Montgomerie, 2013; Loyau et al., 2007; Yoshioka and Kinoshita, 2002). A comparison of cone spectral sensitivity data (Fig 2C) with these reflectance spectra show that the peacock SWS cone is well matched to reflectance from its iridescent peacock blue plumage. While the spectral peaks for the bronze (BZ), blue-green (BG) and outer loose green (GB) barbs agree well with its SWS, MWS and LWS cone spectral ranges, all three also coincide with the same L cone sensitivity for the predator. Both the predator and peafowl VS cone spectral sensitivities also overlap with the reflectance spectrum from the eyespot’s central BB and PB dark violet patches. However, reflectance from these features is weak compared to the other color patches, all of which also reflect weakly in the UV. Fig 2E shows the reflectance spectrum of a representative green leaf, illustrating how its peak at approximately 550 nm and its overall spectra resemble that of the peacock’s loose green barbs; similar spectra for green background foliage have been reported in (Cazetta, Schaefer, and Galetti 2009; Loyau et al. 2007).

Reflectance spectra for parrot feathers (Fig 3E) agreed with previously published values (Tinbergen et al., 2013), confirming a good spectral match between yellow and red feather reflectance with MWS and LWS avian cone sensitivities. The predator L cone sensitivity spans a spectral range corresponding to longer wavelength reflection from both yellow and red pigments (Fig. 3D). Both the yellow and red patches also have reflectance peaks in the UV with a better overlap with bird UVS cone sensitivity than that of VS cones.

A comparison of the peak spectral sensitivities of S and L cones for canids, domestic cats and ferrets with those for the four single cone populations of 21 bird species from 8 different orders (Hart and Hunt, 2007) (Fig 5) illustrates that predator S cones have a peak response similar to that of bird VS, but not UVS, cones. The predator S peak values lie between most peaks of the avian UVS/VS and SWS cone populations, whereas predator M peak values lie between avian MWS and LWS peak values.

**Fig 5.**
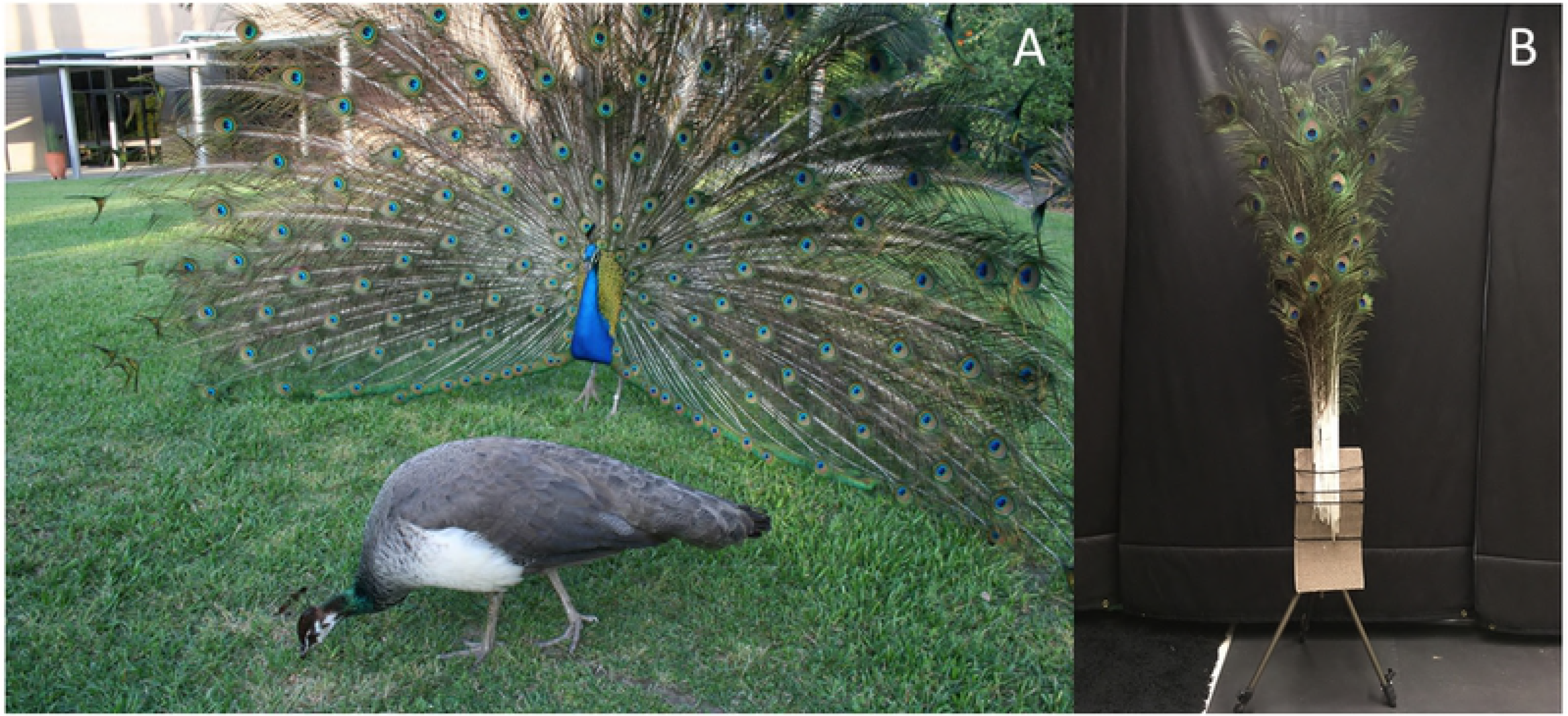
Comparison of peak spectral responses of predator and bird cones. Peak single cone spectral sensitivities for ferret S and L cones (Douglas & Jeffery, 2014) and for bird VS/UVS, SWS,MWS and LWS cones from Fig 5B in (Hart & Hunt, 2007) for 21 species of birds from 9 orders.

### Color and brightness contrast analysis

False color images and analyses using the receptor noise model of visual discrimination are shown in Fig 6–9; all data and PERMANOVA pseudo-F and P values are reported in S1-S4 Datasets. Note that these false color images should be considered as a relative guide and not an absolute indication of the detectability of contrasts because humans have better contrast thresholds than dichromatic mammals by a factor of 3.7 for color and 2 for brightness, as well as similar color contrast thresholds and better brightness thresholds by a factor of ≥ 1.6 for birds (Olsson et al., 2015, 2017). To provide context for the measured color and brightness contrasts between feathers and green saucer magnolia leaves, we also found that the contrasts between leaves from the saucer magnolia and seven other plant species with different shades of green were statistically significant (with one exception out of 27 pairs) with tetrachromatic bird vision *Δ*S_C_ = 2.75 [2.23, 3.28] and *Δ*S_L_ = 3.30 [2.64, 3.97] and dichromatic mammal vision *Δ*S_C_ = 0.55 [0.42, 0.69] and *Δ*S_L_ = 2.53 [2.01, 3.06] (mean [95% CI]) (S3 Fig).

**Fig 6.**
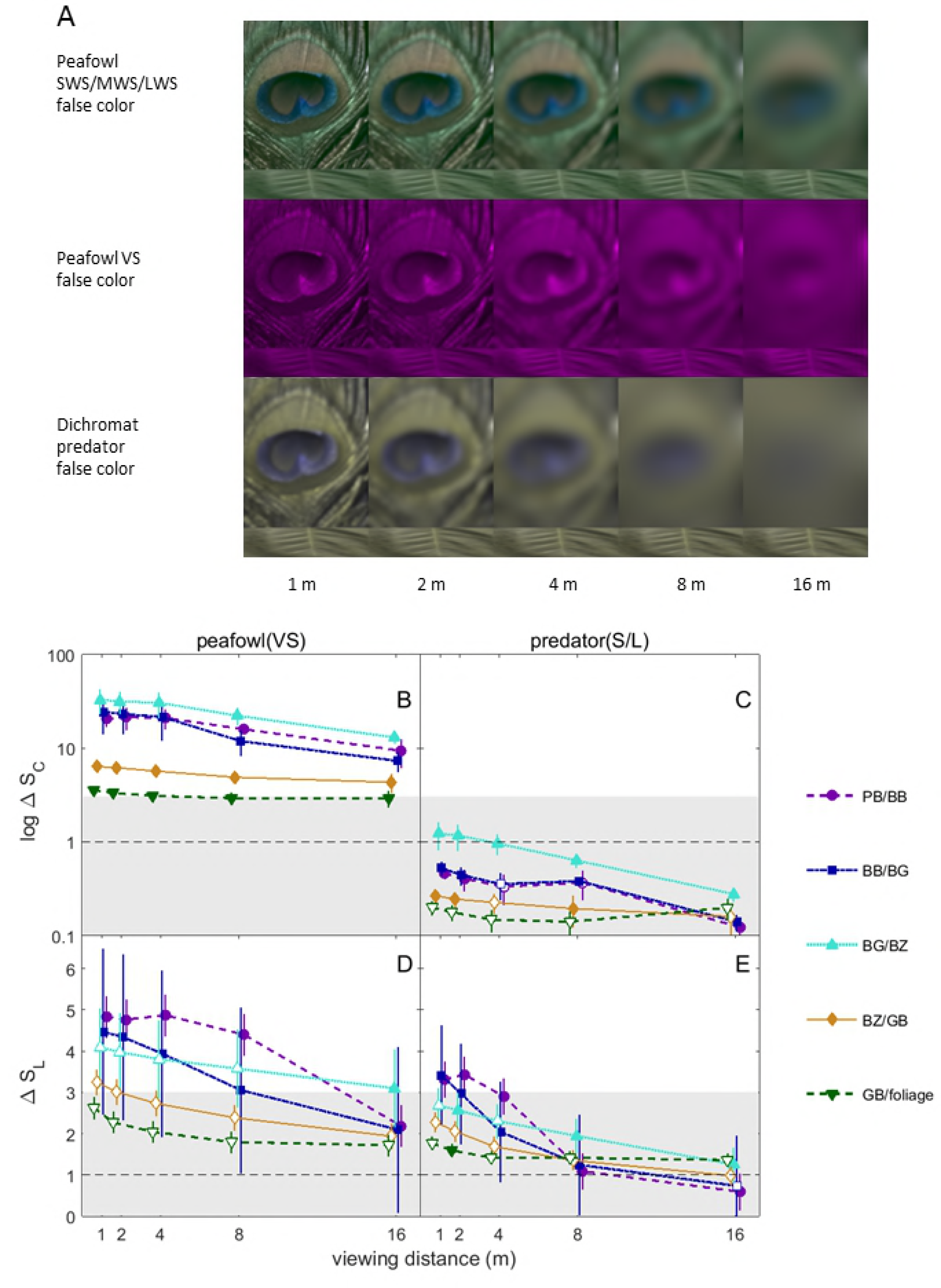
False color images and color and brightness contrast analysis of peacock eyespot feathers. (A) False color images modelling peafowl and dichromatic mammalian predator vision of peafowl eyespot and green leaf (inset at bottom of eyespot image) for different viewing distances. Note that the false color images should be considered as a relative guide and not an absolute indication of the detectability of contrasts because humans have better contrast thresholds by a factor of 3.7 for color and 2 for brightness compared to dichromatic mammals. (B)-(E) Estimated color (delta S_C_) and brightness (delta S_L_) contrasts for adjacent color patches on the peacock eyespots and green vegetation, over a range of viewing distances. All data are shown as grand means with 95% CI error bars. Contrasts corresponding to the same distance have been displaced by horizontal jitter to avoid overlap. Data above the 1 JND line are above the expected threshold for discrimination and contrasts within the grey shaded regions are at most weakly detectable. Closed symbols indicate contrasts that are statistically significant in each organism’s colorspace (i.e., PERMANOVA P < 0.05); note that contrasts that are not statistically significant (closed symbols) due to their large, overlapping variances in the corresponding colorspace may still have mean values greater than the detection threshold.

**Fig 7.**
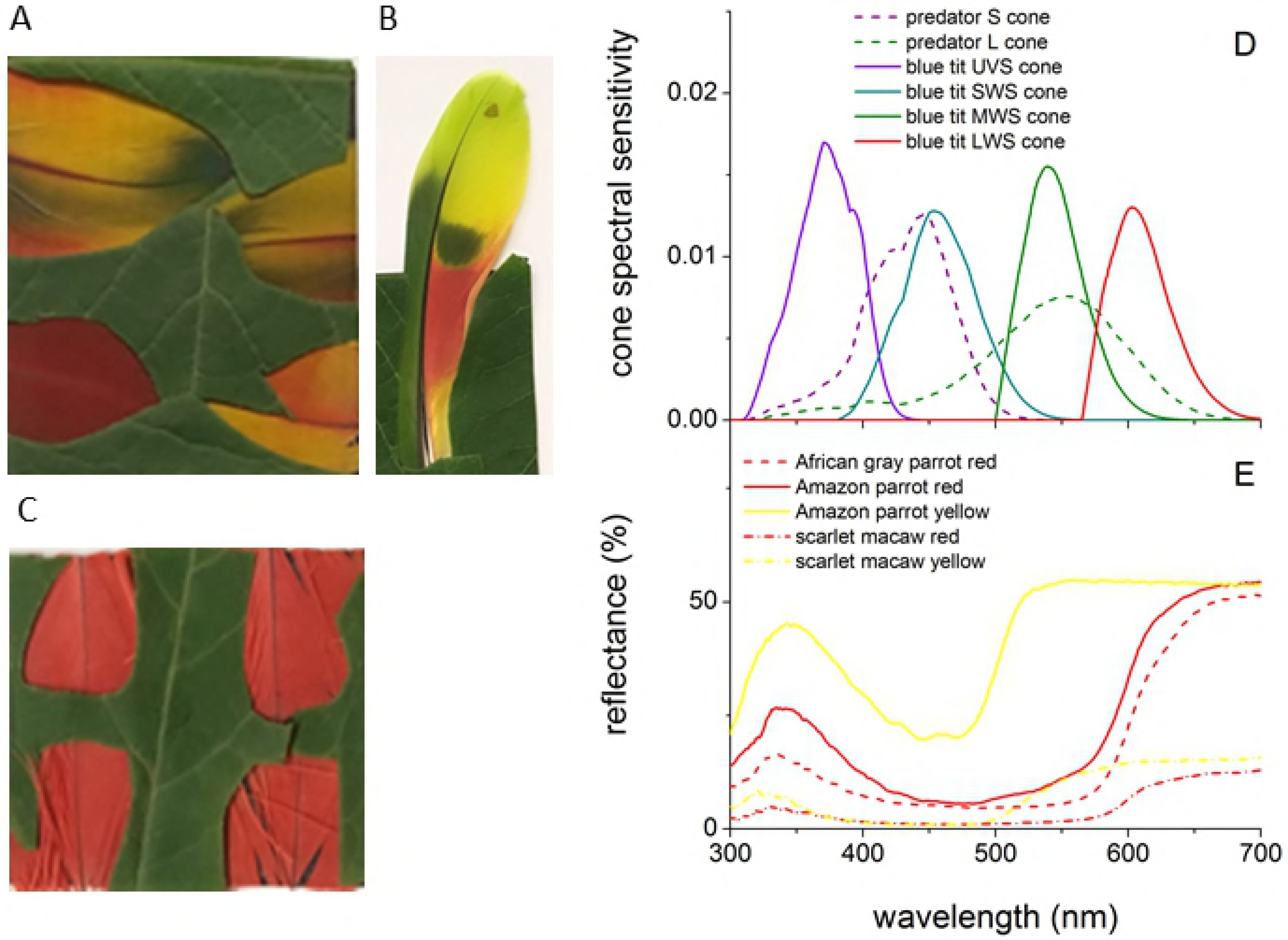
False color images and color and brightness contrast analysis of the peacock model train photographed against various types of vegetation backgrounds. (A) False color images in peafowl and dichromatic mammalian predator vision of peafowl model train for different viewing distances. (B)-(E) Color and luminance contrasts for the model train and features of vegetation, over a range of viewing distances. All data are shown as grand means with 95% CI error bars. See Fig 6 caption for further details.

**Fig 8.**
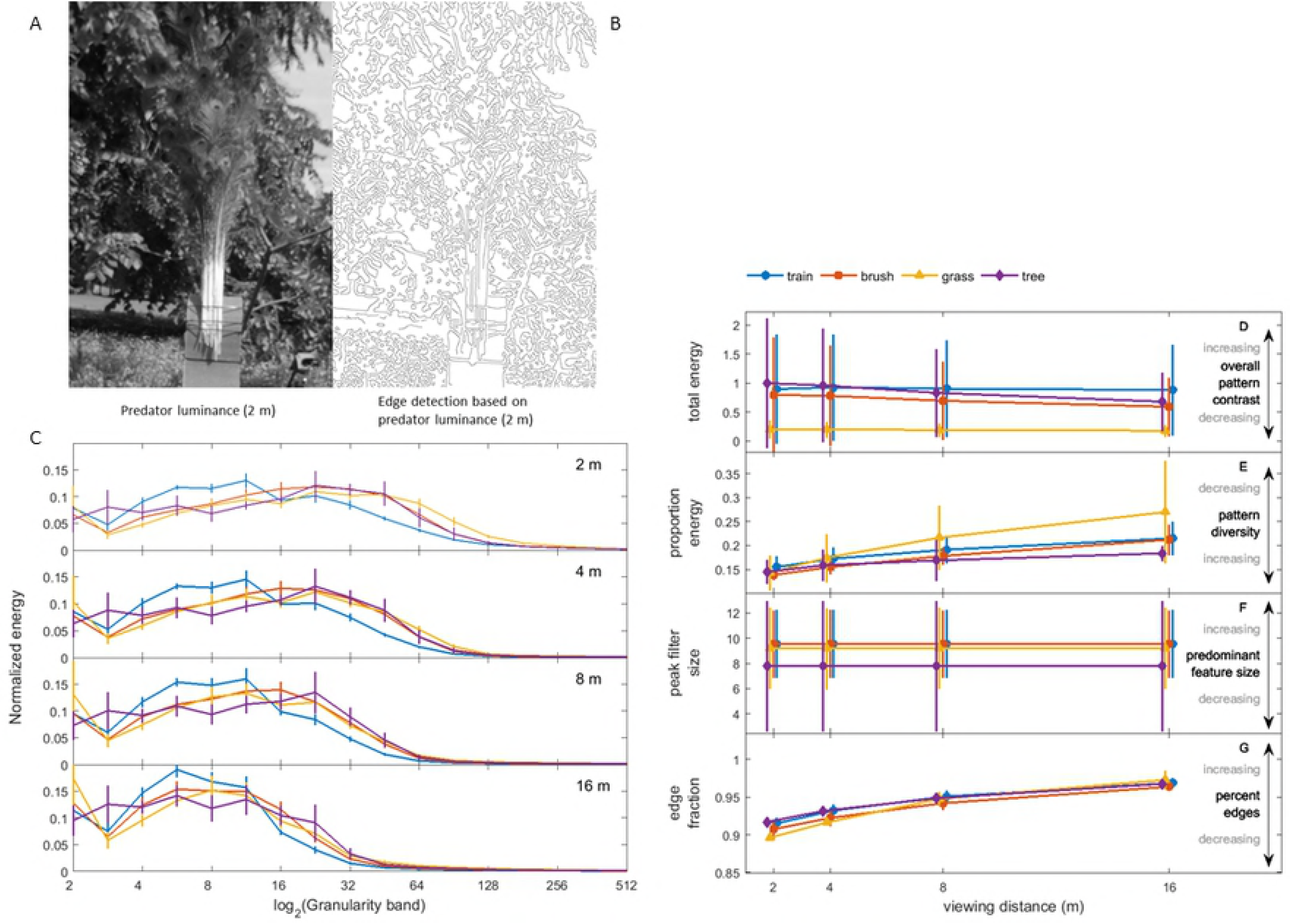
False color images and color and brightness contrast analysis of peacock blue neck feathers used to model the body’s appearance against green foliage. (A) False color images in peafowl and dichromatic mammalian predator vision of peacock blue breast plumage vs green foliage for different viewing distances. (B)-(E) Color and luminance contrasts for the blue plumage relative to green vegetation, over a range of viewing distances. See Fig 6 caption for further details.

**Fig 9.**
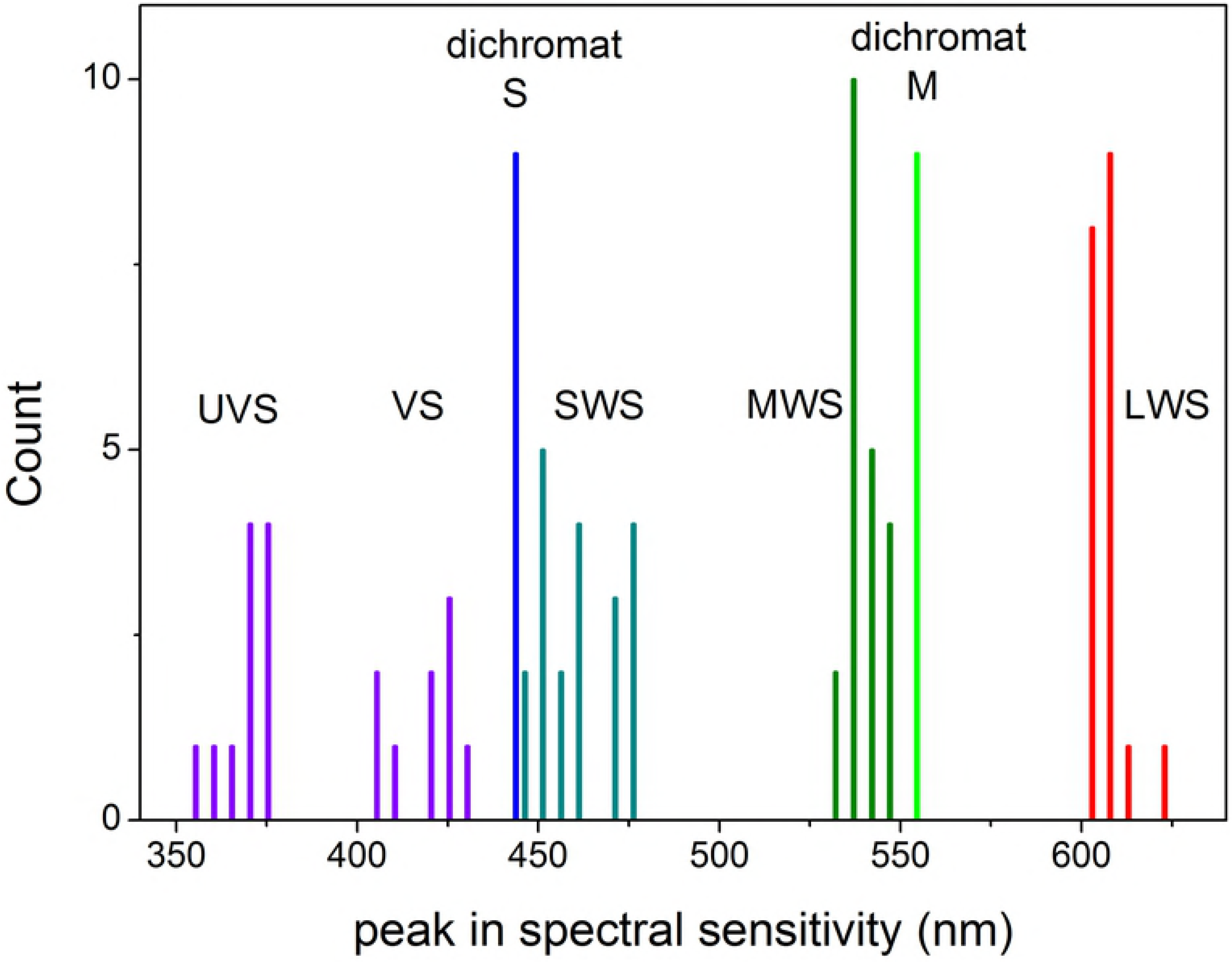
False color images and color and brightness contrast analysis of parrot feathers. (A) False color images in parrot ultraviolet sensitive (UVS) and dichromatic mammalian predator vision of scarlet macaw, African grey parrot and Amazon parrot red and yellow feathers vs green leaf for different viewing distances. (B)-(E) Color and luminance contrasts for parrot feather colors relative to green vegetation, over a range of viewing distances. See Fig 6 caption for further details.

False color images for various simulated viewing distances are displayed for peacock eyespot feathers in Fig 6A and visual signals are plotted vs distance in Fig 6B-E. In peafowl vision, all pairs of adjacent color patches in the peacock’s eyespot give large, statistically significant color contrasts > 3 JND for all distances. The greatest color contrasts were between the blue-green patch and surrounding rings and between the two central pupil-like patches; for some distances ≤ 8 m these same pairs of color patches also had statistically significant brightness contrasts in the 1-3 JND low detectability range. By contrast, in dichromat vision none of the eyespot patch pairs had color contrasts above 1 JND, and only the three innermost pairs of eyespot patches had brightness contrasts that were in the weakly detectable 1-3 JND range.

For peafowl vision, at all distances the model peacock train had statistically significant color and brightness contrasts that were > 3 JND for brush and trees, but not grass (Fig 7; additional false color images in S4 Fig). In dichromat predator vision, all color contrasts for the model train were < 1 JND and brightness contrasts were in the weakly detectable 1-3 JND range. Peacock blue plumage was found to be perceptually detectable by conspecifics at all distances and to lie in the weakly detectable 1-3 JND range for dichromat vision (Fig 8). The false color images of peacock feathers (Fig 6, 7, 8, S4 Fig) demonstrate how color signals relative to background vegetation are diminished when the single dichromat L cone replaces the separate SWS/MWS/LWS cones for birds, especially at larger distances.

Red and yellow parrot feather color patches exhibited large, statistically significant contrasts in avian UVS vision in general, with color contrasts > 3 JND for ≤ 8 m and brightness contrasts > 1 JND for most samples (Fig 9) By contrast, for dichromat vision, none of the red parrot patches and the African grey parrot and scarlet macaw yellow patches had color contrasts > 1 JND, and the Amazon parrot feather yellow patches just exceeded 1 JND for ≤ 4 m. For distances ≤ 8 m, red parrot feather patches had brightness contrasts in the weakly detectable 1-3 JND range for ≤ 8 m and yellow patches had mean values in the range 2.4-6.7 JND.

### Pattern analysis

Fig 10 and S4 Fig show false color images and granularity spectra for the peacock’s train and different types of background vegetation (tall grass, brush and trees) for the various viewing distances modeled (S5 Dataset). Because the spatial frequency of objects in an image as well as an animal’s visual field depend on distance, we would expect objects with similar textures observed at different distances to have similarly-shaped spectra, but possibly different frequency peaks and widths. Granularity spectra for the peacock’s train (Fig. 10B) indeed had the same shape as those for background vegetation in that each had a single broad peak for granularity band values > 3. The peak spatial frequencies of each granularity spectrum moved to lower values as viewing distance increased, as expected from the blurring of fine scale features. For all distances, the peacock’s train and background vegetation had values of proportion energy, peak frequency and total energy that agreed at the 95% CI, with the only exception that the proportion energy for distances > 4 m differed between the model train and trees (Fig. 9D-F). Collectively, these results demonstrate that the peacock’s erect train is an excellent match for the predominant feature size distribution, overall contrast and pattern scale diversity of background vegetation. Moreover, visual examination of the edge detected images (Fig 10A, S5 Fig) suggests that the calculated edge fractions for the train and background foliage agreed at the 95% CI level at all distances (Fig 9G).

**Fig 10.**
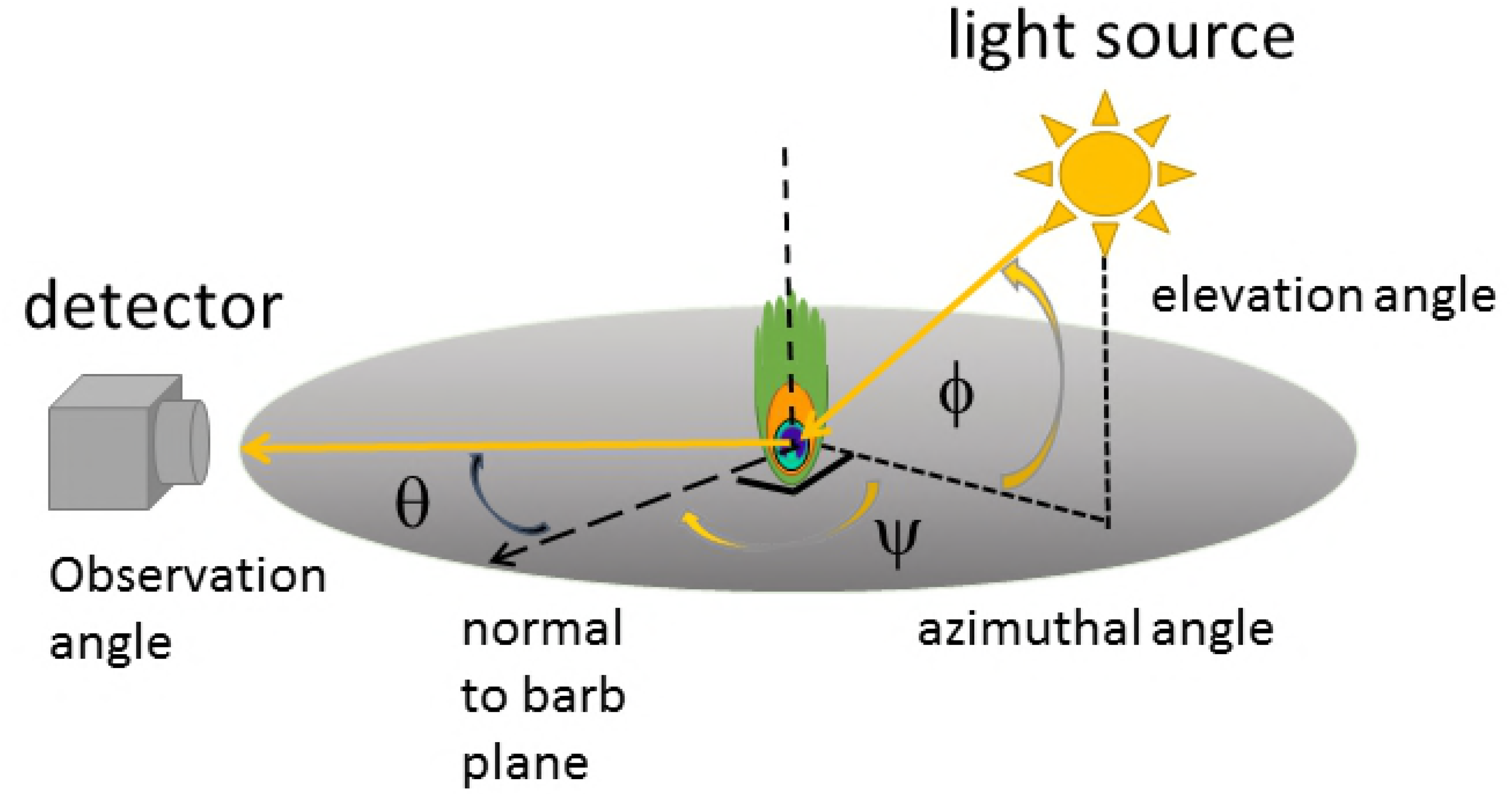
Texture analysis of the model peacock train photographed against various vegetation backgrounds. (A) Image based on dichromatic mammalian predator luminance channel and (B) result of edge detection on the luminance image. (C) Granularity spectrum for the model peacock train and three different regions of vegetation in the background vs viewing distance. (D) Total spectral energy summed over the granularity spectrum, which gives a measure of overall pattern contrast. (E) Proportional energy, a measure of how much the dominant feature size dominates and hence pattern diversity; (F) spatial frequency at peak spectral energy, which is inversely proportional to the predominant feature size; (G) edge fraction, the proportion of the image corresponding to edges. Data are grand means for all model train images and error bars show 95% CI.

## Discussion

The results of our study show that sexually-selected color signals readily detectable by conspecifics are not necessarily conspicuous to mammalian predators. Instead, for all distances considered, the color and brightness contrasts for all feather samples studied here relative to green foliage were much greater for birds than for dichromatic mammals. For viewing distances of 1 m and more, most feather samples (peacock eyespot, model peacock train and red feathers, and all yellow feathers but those of the scarlet macaw at ≤ 2 m) had color contrasts in dichromatic vision that were perceptually indistinguishable from background vegetation at all distances considered here. Unsurprisingly, the same feathers were highly conspicuous to conspecifics: their color contrasts were comparable to values found for avian visual modeling for fruit viewed against green foliage (Cazetta et al., 2009; Fadzly et al., 2013). The brightness contrasts for these feathers vs background foliage in dichromatic predator vision were on the whole greater than the corresponding color signals, although only values for yellow exceeded the weakly detectable 1-3 JND range. This suggests that patterns with high brightness contrast, such as those created by white and dark melanin-pigmented plumage, might be more readily detectable by dichromat predators than color signals, and thus represent a greater detection risk (Montgomerie et al., 2001). While the interpretation of supra-threshold color and brightness contrasts is still debated (Stuart-Fox, 2018), our results show that such supranormal stimuli remain detectable by conspecifics and other birds even at large distances where carnivores cannot perceive them.

Considering parrot feathers in particular, we note that red plumage is at best weakly detectable given its sub-threshold color contrasts and low brightness contrasts when viewed by dichromatic mammals against green foliage, although yellow parrot feathers have brightness contrasts that should be more readily detectable by mammalian predators at close distances. A consideration of cone spectral sensitivities and feather reflectance spectra suggest two reasons for this difference. First, these yellow feathers had an overall higher reflectance than the corresponding red feathers, resulting in their having a higher brightness contrast relative to leaves. Second, yellow feather pigments reflect considerable light in the UV compared to red pigments, whereas UV reflectance is low for green plants. Yellow feathers thus stimulate both predator S and L cones, while green plants primarily stimulate the L cones, providing a mechanism for distinguishing yellow feathers from green foliage backgrounds.

These conclusions should hold for other birds with red and yellow plumage given that a wide variety of species of birds have similar color vision to the species considered here (Fig 5), and that feathers colored with carotenoid pigments have very similar reflectance spectra to the pigment psittacofulvin found in parrot feathers (Shawkey and Hill, 2005; Toral et al., 2008). Thus, our findings indicate that many species of red and yellow feathered birds that appear conspicuous to other birds and humans may in fact be cryptic or poorly visible to predators because of background matching (Stevens and Merilaita, 2011). Our findings also have broader implications for interpreting how color cues, camouflage and possible eye mimicry appear to the majority of mammals. Trichromacy in primates has been suggested to have evolved for a variety of reasons (Carvalho et al., 2017), including detecting ripe fruit and immature leaves, breaking camouflage (e.g., during foraging for eggs) (Troscianko et al., 2017), sexual or social signaling (Hiramatsu et al., 2017), and predator detection (Pessoa et al., 2014). Our results suggest that the evolution of trichromacy may also have provided catarryne primates, howler monkeys and some marsupials with an advantage in detecting colorful birds, reptiles, amphibians and insects.

Ultraviolet vision per se does not result in these differing visual signals: these dichromatic mammalian predators have similar near-UV S cone spectral sensitivity to the VS cones of birds (Douglas and Jeffery, 2014; Stevens and Cuthill, 2007). Indeed, as noted above, since red and yellow parrot feathers and the central patches on peacock feathers reflect appreciable UV light this may make these feathers more detectable by dichromats. It is therefore important to include UV reflectance in modeling of visual signals in dichromatic mammalian visual systems, as opposed to relying on image processing of human visible RGB photographs (Pongrácz et al. 2017). These results also are not merely a consequence of birds having more types of cones than carnivores: given the similar spectral response of dichromatic mammal S and L cones and avian VS and MWS cones, color patches could in principle generate similar contrasts in both visual systems. Instead, these feathers have low contrast in dichromatic mammal vision due to a combination of low visual acuity, higher receptor noise levels and poorer spectral discrimination over the L cone response range.

Focusing now on peacock eyespots, the large color contrasts for peafowl vision arise from spectral tuning between the reflectance spectra of each peacock eyespot color patch and peafowl single cone spectral sensitivities, similar to the agreement reported earlier between red and yellow pigment reflectance spectra and tetrachromatic UVS cone responses for parrot plumage and vision (Tinbergen et al., 2013). It is especially notable that the greatest color contrasts are due to the blue-green ring, since its iridescence has been found to correlate with peacock mating success (Dakin and Montgomerie, 2013; Loyau et al., 2007), and its chromatic contrast was calculated to be the most salient signal in images of a displaying peacock (Pike, 2018).

When we computed measures for the model peacock train against a foliage background in dichromatic predator vision, the train feathers were found to have below detection threshold color contrasts and brightness contrasts in the low detectability 1-3 JND range, similar in magnitude to those for various types of green vegetation. Taken together with the eyespot and textural analysis results, this indicates that dichromatic mammalian predators are likely unable to discriminate the peacock’s train from green vegetation during foraging, although the eyespot’s innermost features create low detectable brightness contrasts at nearby distances. Contrary to common assumption, this suggests the counterintuitive hypothesis that that both the color and brightness contrasts might function as disruptive camouflage for peacocks viewed from a distance, making the train difficult to distinguish as a whole (Hillgarth, 1984; Ridley et al., 1984; Stevens and Merilaita, 2011).

The peacock’s blue plumage had large, detectable levels of color contrast at all distances for peafowl vision, though both color and brightness contrasts were in the low detectable range for dichromatic predator vision. Thus, the blue head, neck and breast contour feathers may represent a greater visual signal for distant conspecifics, as well as a greater predation risk, than the much larger train; however, all of these values are likely less conspicuous when forest shade diminishes their blue hue. Given that noniridescent blue feathers from other birds have been shown to have similar reflectance spectra to peafowl blue plumage, these results are likely generalizable (Osorio and Ham, 2002).

Do peacock eyespots mimic eyes in appearance when viewed by conspecifics or by predators? Eyes and eye-like patterns have been shown to be highly salient visual signals for birds, humans and domestic dogs (Somppi et al., 2016; Stevens and Ruxton, 2014; Yorzinski et al., 2015). However, our false color images show that peafowl eyespots do not always appear to have a central dark, circular pupil when viewed at typical display distances, either in peafowl or dichromatic mammalian predator visual models (Fig 11A,B,C). Therefore, it is not obvious that peacock ocelli appear eye-like to nearby viewers. On the other hand, blurring at larger distances ≤ 8 m causes eyespots to appear to have pupil-like dark centers in dichromatic predator vision, indicating that they may indeed appear eyelike to predators when viewed at this intermediate range (Fig 11D).

**Fig. 11.**
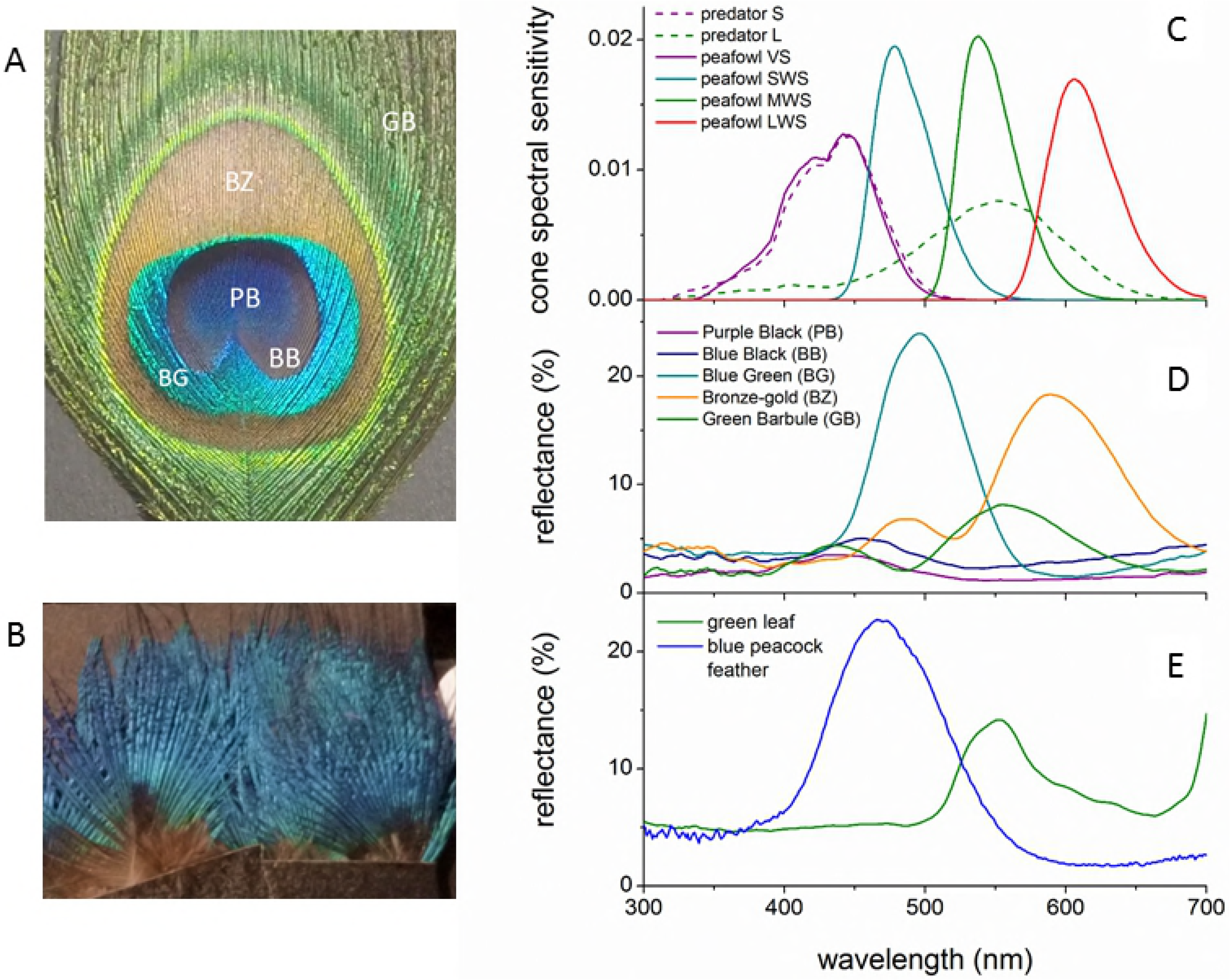
False color images of peacock eyespots show that eye-mimicry depends on viewing distance. At 1 m, peacock eyespots have central features with brightness comparable to the surrounding rings in false color images for peafowl vision in the (A) LWS/MWS/SWS and (B) VS cone channels, and for dichromatic mammals (C). False color image of the model train in dichromatic mammalian vision at 2 meters (D) show that the eyespots feature a darker pupil-like center at distances ≤ 8 m.

Taken as a whole, these results call into question the widely-held assumption that the peacock’s eyespot feathers are highly conspicuous for all viewers. Indeed, (Hillgarth, 1984) noted that even to humans the peacock’s train can be well camouflaged against foliage in its native habitats and (Ridley et al., 1984) has suggested that peafowl eyespots originated as a form of camouflage for their native dappled light environments. Motion cues during peafowl displays and other behaviors might enhance the detectability of their visual signals, although evidence is mixed whether motion increases or decreases visual contrast thresholds (Hodos et al., 2003). On the other hand, the motions of the flexible loose green barbs in the train might also simulate that of background brush and grasses, a visual illusion studied in insects, crabs, spiders and lizards but not yet in birds (Peters et al., 2008). Future video studies could utilize cameras adapted for multispectral imaging provided with a filter that transmits light from the near-UV to 700 nm.

Supporting these conclusions, we have been unable to find any data in the literature supporting the idea that the peacock with trains experience a significant, let alone an enhanced, risk of predation compared to, e.g., either peacocks without trains or peahens. In fact, one survey of a feral peafowl population (Petrie, 1992) found that predated adult peacocks tended to have relatively small trains and lower mating success. Another study found that peafowl are preyed on far less by leopards than expected given that they were the most abundant prey species in the region studied (Mondal et al., 2011). Adult peacocks are reported to have several effective antipredator strategies, including running (Ali and Ripley, 1981; Thavarajah et al., 2016; Wilkinson et al., 2015), flight (Askew, 2014), fighting with their sharp spurs (Petrie et al., 1991), hiding in dense thickets (Ali and Ripley, 1981; Harikrishnan et al., 2010; de Silva et al., 1996), and using group vigilance along with alarm calling (Yasmin and Yahya, 2000). At dusk, they roost in high trees chosen for their protection against predators (de Silva et al., 1996; Trivedi and Johnsingh, 1996). As a result, peafowl appear to be at highest risk of predation primarily during their first year (de Silva et al., 1996).

The available evidence thus indicates that any putative enhanced risk of predation or other handicap suffered by adult peacocks is likely to be incurred by factors other than visual signals created by their eyespot train feathers. For example, peacocks spend a large percentage of their time maintaining their plumage (Walther, 2003) and displaying (Dakin and Montgomerie, 2009; Harikrishnan et al., 2010). Thus, the elaborate courtship displays of peacocks may correspond to handicaps due to the conspicuous appearance of their blue plumage, time lost from foraging for food due to plumage maintenance and courtship displays, the male’s likely inattention to predators during displays, and the metabolic demands of the male’s courtship displays (Vehrencamp et al., 1989).

It is important to note that several other factors may result in feathers having even lower visual contrasts in predator vision than reported here. First, our viewing and illumination geometries were chosen to optimize color and brightness cues. Second, when feathers are viewed at low illumination levels either in dim forest lighting or at twilight or night, they also are likely to have lower color and brightness contrasts (Freitag and Pessoa, 2012; Sicsú et al., 2013). Furthermore, since UV reflectance helps distinguish feathers from green foliage, the reduction in UV irradiance in forest shade is likely to render feathers less detectable in forest shade than in direct sunlight (Endler, 1993). Third, we also modeled only distance-dependent blurring due to visual acuity (retinal sampling), but a more complete treatment would use each species’ behaviorally measured contrast sensitivity function (CSF) (Melin et al., 2016) to account for the optics of the eye and other factors that have determined for our study species (Jarvis and Wathes, 2007). While we lacked the data to perform this additional analysis, the additional blurring would make dichromatic predators even less likely to be able to detect the feathers than our estimated contrasts indicate. Fourth, we chose to compare feathers with relatively dark green leaves (S3 Fig). The small variation in color contrasts indicates our findings are generalizable to other shades of green, while suggesting that feathers should be even less visible against light green foliage. Thus, our results likely overestimate the detectability of these feathers by dichromatic mammals.

The largest source of uncertainty in our analysis is the lack of behaviorally-measured Weber fractions for color contrast for terrestrial carnivorans for the conditions considered here. This data also would be valuable for studies that often have had to rely on human visual modeling in analyzing egg camouflage (Stoddard et al., 2016) and the relationship between plumage, brightness and antipredator vigilance (Pascual et al., 2014). The most relevant measures would involve behavioral tests to determine whether these mammals can detect feathered model birds when other cues (e.g., olfactory) are controlled for. Any such studies ought to be sure to use illumination sources that include UV, as well as color cues that closely match the reflectance spectra of natural objects (Stoddard et al., 2019).

Darwin stated that “Even the bright colors of many male birds cannot fail to make them conspicuous to their enemies of all kinds” (Darwin, 1888). On the contrary, our study implies that some species of birds that appear vividly colorful to humans and other birds may appear drab and inconspicuous in the eyes of mammalian predators. This conclusion is consistent with the finding that susceptibility to cat predation does not correlate significantly with sexual dichromatism in birds (Møller et al., 2010). The predation risk incurred by colorful plumage instead depends on specifics of pigmentation, photoreceptor response, and environmental context, as suggested by sensory drive theory (Cummings and Endler, 2018), and thus should be assessed on a case-by-case basis.

Predators have a variety of other means of detecting prey, including visual motion perception and sensing acoustic, tactile and olfactory cues. Our results highlight the importance of understanding how dynamic behaviors during multimodal displays, foraging and other activities make birds more apparent to mammals and other predators than do their seemingly-conspicuous colors alone.

## Acknowledgements

We wish to thank Cassie Stoddard for providing us with her edge detection code, Charles Chubb and Roger Hanlon for their granularity code, and Robert Beyer for instrumentation design.

## Author contributions

Conceptualization: S.A.K., R.D.; Methodology: S.A.K., R.F., Y.L., Y.W; Investigation: S.A.K, R.F., Y.L., Y.W; Data curation: S.A.K., Y.W.; Software: S.A.K., Y.W.; Validation: S.A.K., Y.W.; Visualization: S.A.K., R.F., Y.L., Y.W; Formal analysis: S.A.K.,.W.; Writing - original draft: S.A.K.; Writing - review & editing: S.A.K., R.D., R.F., Y.L., Y.W.

## Supporting information captions

S1 Appendix. Predators of Indian peafowl.

S2 Appendix. Species of plants used for background foliage in multispectral images.

S3 Appendix. Methods for spatially filtering multispectral images to account for visual acuity effects at varying viewing distances.

S1 Fig. Multispectral camera specifications: image sensor spectral response and filter transmission spectra.

S2 Fig. Reflectance spectroscopy apparatus and multispectral camera filming setup.

S3 Fig. Various green leaves imaged and analyzed for comparison with feather samples and the saucer magnolia leaves used as a background.

S4 Fig. Model train false color images.

S5 Fig. Model peacock train edge detected images.

S1 Dataset. Color and brightness contrast data and PERMANOVA pseudo-F and P values for Indian peacock eyespot feathers

S2 Dataset. Color and brightness contrast data and PERMANOVA pseudo-F and P values for model Indian peacock train

S3 Dataset. Color and brightness contrast data and PERMANOVA pseudo-F and P values for Indian peacock blue body feathers

S4 Dataset. Color and brightness contrast data and PERMANOVA pseudo-F and P values for parrot feathers

S5 Dataset. Granularity and edge-detection visual texture data for model Indian peacock train

